# Dynamic expressions of confidence within an evidence accumulation framework

**DOI:** 10.1101/2020.02.18.953778

**Authors:** Kobe Desender, Tobias H. Donner, Tom Verguts

## Abstract

Human observers can reliably report their confidence in the choices they make. An influential framework conceptualizes decision confidence as the probability of a decision being correct, given the choice made and the evidence on which it was based. This framework accounts for three diagnostic signatures of human confidence reports, including an opposite dependence of confidence on evidence strength for correct and error trials. However, the framework does not account for the temporal evolution of these signatures, because it only describes the transformation of a static representation of evidence into choice and the associated confidence. Here, we combine this framework with another influential framework: dynamic accumulation of evidence over time, and build on the notion that confidence reflects the probability of being correct, given the choice and accumulated evidence *up until that point*. Critically, we show that such a dynamic model predicts that the diagnostic signatures of confidence depend on time; most critically, it predicts a stronger opposite dependence of confidence on evidence strength and choice correctness as a function of time. We tested, and confirmed, these predictions in human behaviour during random dot motion discrimination, in which confidence judgments were queried at different points in time. We conclude that human confidence reports reflect the dynamics of the probability of being correct given the accumulated evidence and choice.

## Introduction

Human observers can provide very precise judgments about the confidence of their choices. They often claim high confidence for correct choices and low confidence for errors. Accurate confidence estimates are adaptive because confidence is used to regulate future behaviour (Desender, Boldt, et al., 2019; Desender, Murphy, et al., 2019; van den Berg et al., 2016). An influential framework posits that internal representations of decision confidence, and agents’ overt reports thereof, reflect the probability of being correct, given the choice made and given the evidence on which it was based (Kepecs et al., 2008; Pouget et al., 2016; Sanders et al., 2016). In this framework, both choice and confidence are directly based on the same underlying computations. This principled approach predicts three key qualitative signatures of confidence (Kepecs et al., 2008): Signature 1: an interaction between evidence strength and choice accuracy, whereby confidence increases with evidence strength for correct choices, but decreases for incorrect choices; Signature 2: confidence increases monotonically with accuracy; Signature 3: a steeper increase in performance for high versus low confidence trials as a function of evidence strength. These three signatures have been observed in neural data (Kepecs et al., 2008), several implicit behavioural measures of confidence (Braun et al., 2018; Kepecs et al., 2008; Sanders et al., 2016; Urai et al., 2017), and explicit confidence reports of human observers (Fleming et al., 2018; Sanders et al., 2016). An important limitation of this framework is that it is static: a fixed quantity of evidence determines both the choice and associated confidence. Therefore, this framework does not account for the dynamics of decision-making, the associated trade-off between speed and accuracy, and their effect on confidence reports.

To address this issue, we combine this framework with an evidence accumulation model. According to evidence accumulation models, perceptual decisions are based on a gradual accumulation of noisy sensory evidence over time (Gold & Shadlen, 2007; Ratcliff & McKoon, 2008). In two-choice tasks, a decision maker accumulates evidence for each option, and the option for which the integrated evidence first crosses a decision threshold is selected, indicating commitment to choice (Ratcliff & McKoon, 2008). The efficiency (i.e., signal-to-noise ratio) of the accumulation process is governed by the so-called drift rate. To account for confidence within such a dynamic model, Kiani and Shadlen (2009) proposed that confidence reflects the probability of being correct, given choice, evidence *and time*.Although such conceptualization explains the scaling of confidence with evidence strength (Kiani & Shadlen, 2009), with decision time (Kiani et al., 2014) and with evidence volatility (Zylberberg et al., 2016), an often cited limitation of this account is that it cannot explain why human confidence judgments are higher for correct than for error trials within a single level of evidence strength (Pleskac & Busemeyer, 2010). Put differently, this model cannot easily explain how humans become aware of their mistakes. In order to account for this, it has been proposed that evidence continues to accumulate after the decision boundary has been reached (Moran et al., 2015; Pleskac & Busemeyer, 2010; Resulaj et al., 2009; Van Den Berg et al., 2016; Yu et al., 2015). Confidence then reflects (a transformation of) the accumulated evidence after this period of post-decision accumulation. With non-zero drift rate, such a model will on average accumulate further evidence in favor of the selected choice when it was correct, whereas it will on average accumulate evidence for the non-selected choice in case of an error (i.e., potentially leading to a change of mind; Van Den Berg et al., 2016). Thus, the process of post-decision evidence accumulation allows the model to detect its own errors in a graded fashion. Such dynamic models quantifying confidence as a function post-decisional evidence and decision time have been fruitful in explaining decision confidence (Calder-Travis et al., 2020; J. Drugowitsch et al., 2012; Jan Drugowitsch et al., 2014; Moran et al., 2015; Moreno-Bote, 2010; Pereira et al., 2020; Pleskac & Busemeyer, 2010; Van Den Berg et al., 2016; Yu et al., 2015).

It remains unclear, however, whether such a model also accounts for the three diagnostic signatures of confidence identified by Sanders et al. (2016), and if so how these signatures depend on the duration of post-decision accumulation. Here, we address this question through the combination of simulations of a popular evidence accumulation model, the drift diffusion model (DDM), and the analysis of human behavior in a perceptual decision task. First, we show that these signatures of confidence dynamically depend on post-decision processing time, a pattern that was also seen in human participants. Second, if confidence is quantified after a period of post-decision accumulation, a stopping criterion is needed, similar to the decision boundary for first-order choices. Here, using a manipulation of evidence volatility, we shed light on the question whether human participants use a time-based stopping rule (i.e., terminate sampling after a certain amount of time) or an evidence-based stopping rule (i.e., terminate sampling after the accumulated evidence reached a certain threshold value).

## Methods

### Participants

Thirty participants (two men; age: *M* = 18.5, *SD* = .78, range 18 – 21) took part in return for course credit. All participants reported normal or corrected-to-normal vision and were naïve with respect to the hypothesis. All but four participants were right handed. Four participants were excluded because their performance was not different from chance level in the immediate condition (as assessed by a binomial test). All data have been made publicly available via the Open Science Framework and can be accessed at osf.io/83×7c. Non-overlapping analyses of these data have been published elsewhere (Desender, Boldt, et al., 2019).

### Ethics Statement

Participants provided written informed consent before participation. All procedures were approved by the local ethics committees at Ghent University.

### Stimuli and apparatus

Stimuli were presented in white on a black background on a 20-inch LCD monitor with a 75 Hz refresh rate, using Psychtoolbox3 (Brainard, 1997) for MATLAB (The MathWorks, Natick, MA). Random moving white dots were drawn in a circular aperture centered on the fixation point. The current experiment was based on code provided by Kiani and colleagues (2013). Parameter details can be found there.

### Procedure

Participants were presented with dynamic random dot motion and were asked on each trial to decide as fast and accurate as possible whether a subset of dots was coherently moving towards the left or the right side of the screen (See Figure 1). Each experimental trial started with a fixation dot for 750ms followed by random dot motion that lasted until a response was made, with a maximum of 3 seconds. On each trial, the difficulty of these decisions was controlled by manipulating the percentage of coherently moving dots, which was either 0%, 5%, 10%, 20% or 40%. In each block, there was an equal number of leftward and rightward movement, and the different coherence levels were randomly intermixed within a block. We also manipulated the volatility of motion coherence over the course of a single trial. In the low evidence volatility condition, the proportion of coherently moving dots was the same on every timeframe within a trial. In the high evidence volatility condition, the proportion of coherently moving dots was on each timeframe sampled from a Gaussian distribution with mean equal to the generative distribution of that trial and a standard deviation of .256. In the high volatility condition, additional noise is thus introduced in the decision process. Previous work has shown that this manipulation, which speeds up RTs and increases confidence, is diagnostic for an evidence-based stopping rule for immediate decisions (Zylberberg et al., 2016). We here used this manipulation to shed light on the stopping rule for delayed confidence reports.

**Figure 1.**
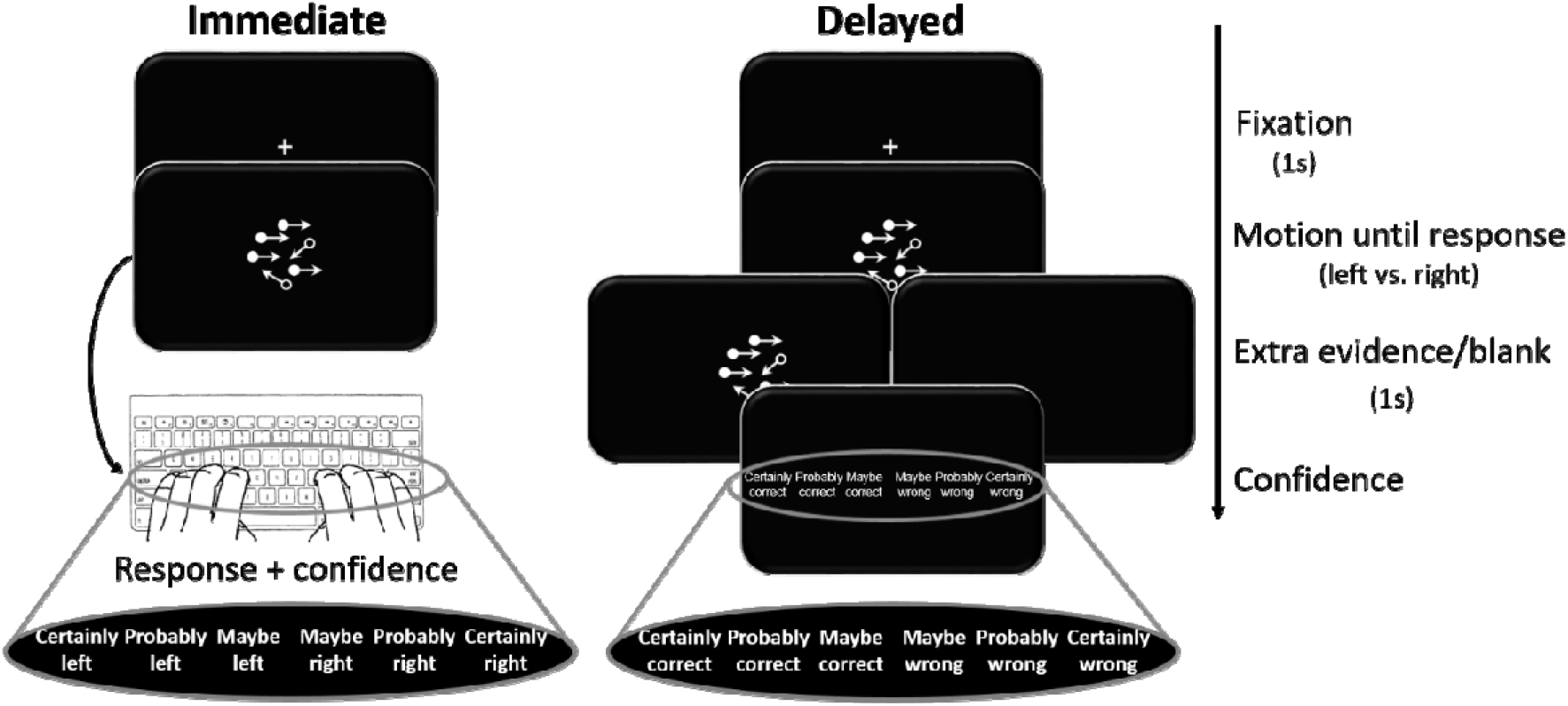
Experimental task. Sequence of events in the experimental task. Participants decided, as fast and accurately as possible, whether the majority of dots were moving left or right. In the immediate condition, they did so by jointly indicating their choice (left or right) and confidence (sure correct, probably correct or guess correct) with a single response. In the delayed condition, participants first indicated their choice with their thumbs (left or right), and after a 1s blank or 1s of continued motion, they indicated the degree of confidence in their decision using a six-point confidence scale (ranging from certainly correct to certainly wrong).

In order to query the dynamics of post-decision processing and its influence on the diagnostic signatures of confidence there were three different interrogation conditions. In the *immediate* condition, participants jointly indicated their response and their level of confidence. The numerical keys ‘1’, ‘2’, ‘3’, ‘8’, ‘9’, and ‘0’ on top of the keyboard mapped onto ‘sure left’, ‘probably left’, ‘guess left’, ‘guess right’, ‘probably right’, and ‘sure right’, respectively. In the *delayed blank* condition, participants indicated their response (left or right) by pressing ‘c’ or ‘n’ with the thumbs of their right and left hand, respectively. Then, a blank screen was presented for 1s, after which the following six confidence options were presented on the screen: ‘sure correct’, ‘probably correct’, ‘guess correct’, ‘guess error’, ‘probably error’, ‘sure error’ (reversed order for half of the participants). Participants had unlimited time to indicate their level of confidence by pressing one of the corresponding numerical keys (i.e., ‘1’, ‘2’, ‘3’, ‘8’, ‘9’, and ‘0’) on top of the keyboard. The *delayed extra evidence* condition was similar to the delayed blank condition, except that now 1s of continued random motion was presented during the 1s interval between the response and the confidence judgment. The continued motion had the same direction, the same motion coherence and the same level of evidence volatility as the pre-decisional motion.

The entire experiment comprised twelve blocks of sixty trials each, including three practice blocks. In the first practice block, participants only indicated the direction of the dots (i.e., no confidence), and each trial stopped after a response was given. Only coherence levels of .2 and .4 were presented. When participants made an error, the message ‘Error’ was shown on the screen for 750ms. This block was repeated until mean accuracy exceeded 75%. The second practice block was similar, except that now the full range of coherence levels was used. This block was repeated until mean accuracy exceeded 60%. Block three served as a last practice block, and was identical to the main experiment. No more feedback was presented from this block on. Each participant then performed three blocks of each interrogation condition, with the specific order depending on a Latin square. Before the start of block seven and block ten (i.e., start of a new interrogation condition), participants performed eight practice trials with .4 coherence using the procedure of the subsequent block, to get familiarized with the response keys. These eight trials were repeated until accuracy exceeded 75%. After each block, participants received feedback about their performance in that block, including mean response time on correct trials, mean accuracy, and the absolute value of the correlation between accuracy and confidence. Participants were motivated to maximize these three values.

### Statistical analyses

Behavioral data and model predictions were analyzed using mixed regression modeling. This method allows analyzing data at the single-trial level. We fitted random intercepts for each participant; error variance caused by between-subject differences was accounted for by adding random slopes to the model. The latter was done only when this significantly increased the model fit. RTs and confidence were analyzed using linear mixed models, for which *F* statistics are reported and the degrees of freedom were estimated by Satterthwaite’s approximation (Kuznetsova et al., 2014). Accuracy was analyzed using logistic linear mixed models, for which *X*^*2*^ statistics are reported. Model fitting was done in R (R Development Core Team, 2008) using the lme4 package (Bates et al., 2015).

### Drift diffusion model

#### Fitting

Drift diffusion model parameters were estimated using hierarchical Bayesian estimation within the HDDM toolbox (Wiecki et al., 2013). The HDDM uses Markov-chain Monte Carlo (MCMC) sampling, which generates full posterior distributions over parameter estimates, quantifying not only the most likely parameter value but also uncertainty associated with each estimate. Due to the hierarchical nature of the HDDM, estimates for individual subjects are constrained by group-level prior distributions. In practice, this results in more stable estimates for individual subjects. For each model, we drew 100.000 samples from the posterior distribution. The first ten percent of these samples were discarded as burn-in and every second sample was discarded for thinning, reducing autocorrelation in the chains. Group level chains were visually inspected to ensure convergence, i.e. ruling out sudden jumps in the posterior and ruling out autocorrelation. Additionally, all models were fitted three times, in order to compute the Gelman-Rubin R hat statistics (comparing within-chain and between-chain variance). We checked and confirmed that all group-level parameters had an R hat between 0.98-1.02, showing convergence between these three instantiations of the same model. Because individual parameter estimates are constrained by group-level priors, data for different subjects are dependent, and frequentist statistics cannot be used. The probability that a condition differs from another can be computed by calculating the overlap in posterior distributions.

When fitting the data (choices and reaction times), both drift rate (v) and decision bound (a) were allowed to vary as a function of coherence and evidence volatility, whereas non-decision time (ter) was fixed across conditions. According to our hypothesis, the effect of evidence volatility will be expressed in the within-trial variability parameter (σ). When fitting the DDM this parameter is fixed (i.e., to .1 in the Ratcliff Diffusion model or to 1 in the currently used HDDM). Because σ is a scaling factor, after fitting the model, we next scaled drift rate, decision bound and within-trial variability for each condition so that decision bound was equal to 1. Thus, this approach allows estimating within-trial variability. Note that under this approach, an implicit assumption is that the decision bound does not differ between the different conditions.

#### Simulations

Using the estimates obtained from the HDDM fit, predictions were generated using a random walk approximation of the diffusion process (Tuerlinckx et al., 2001). This method simulates a random walk process that starts at z*a (here, *z* was an unbiased starting point of .5) and stops once the integrated evidence crosses 0 or a. At each time interval *t*, a displacement Δ occurs with probability *p* and a displacement -Δ with probability *1-p*. Both quantities are given in equation (1).

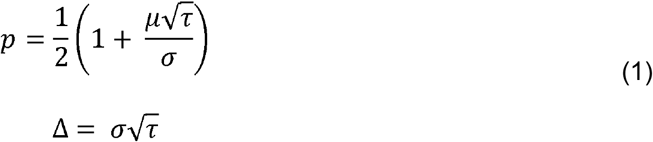

Drift rate is given by *μ*, and within-trial variability is given by *σ*. In all simulations *τ* was set to 1e-4. In order to construct the heat map representing the probability of being correct shown in Figure 2, 300.000 random walks without absorbing bounds were generated, using the fitted drift drates. This assured sufficient data points across the relevant part of the heat map. Subsequently, the average accuracy was calculated for each (response time, evidence) combination, based on all trials that had a data point for that (response time, evidence) combination. Smoothing was achieved by aggregating over evidence windows of .01 and *τ* windows of 2.

**Figure 2.**
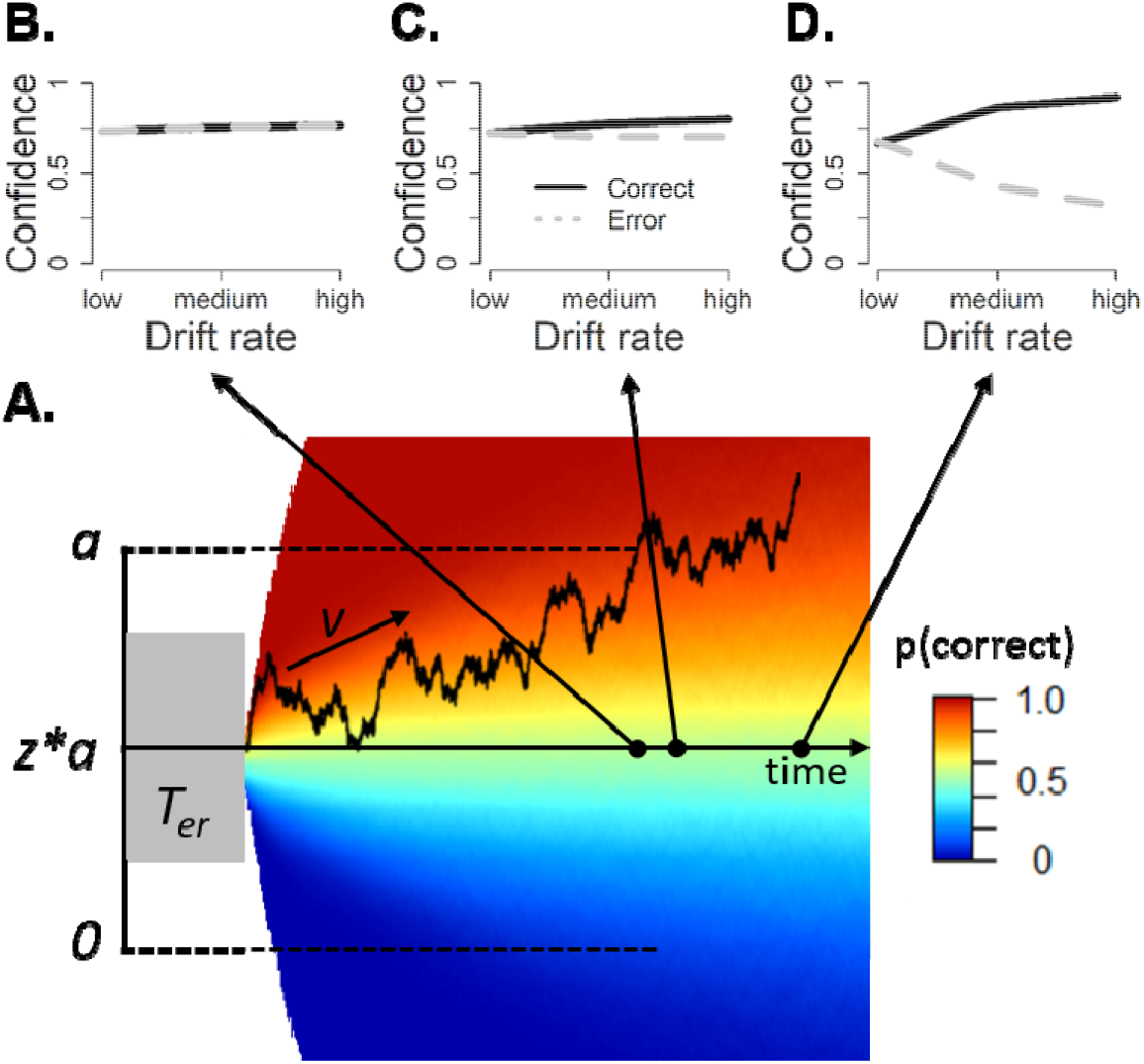
Quantifying decision confidence within an evidence accumulation framework. **A**. Noisy sensory evidence is accumulated over time, until the decision variable reaches one of two bounds (a or 0), corresponding to a left or right choice, respectively. After the decision variable reaches a bound, evidence continues to accumulate. The heat map shows the probability of being correct given time and evidence, conditional on reaching the upper boundary (i.e., making an “up” response). Confidence is quantified as the probability of the choice being correct, given elapsed time and the integrated evidence (i.e., represented by the color of the heat map). Confidence can be queried at different points in time. **B-D**. Model predictions about signature 1, an interaction between evidence strength and accuracy, depending on whether confidence is quantified at the same time as the choice (B; i.e. at boundary crossing), after a small temporal delay (C), or after considerable additional post-decision accumulation (D).

To generate model fits for choices and RTs and model predictions for confidence, we used the parameters obtained by the HDDM fit. For each combination of coherence levels, within-trial evidence volatility and interrogation condition, we simulated 5000 trials per participant. Both immediate and delayed confidence predictions were obtained by reading out the probability of being correct from the heat map given RT and evidence, conditional on the response given. Model predictions about confidence were then converted from a continuous scale to a categorical scaling by dividing them into three (immediate condition) or six (delayed condition) equal-sized bins. For the immediate condition, confidence predictions were obtained without any post-decision accumulation. In the adapted version confidence was quantified with a small temporal delay of .1s; other (small) values led to very similar results. For delayed confidence predictions with a time-based stopping rule, after reaching the decision bound, the random walk process continued for one second (i.e., the duration of the ITI) plus the average response speed of confidence judgments in that condition minus the non-decision time of that condition. For the evidence-based stopping rule, after the evidence crosses *a*, an evidence-based stopping rule (i.e., a horizontal boundary) was placed at *a + a\**.*125* and 0 (or similarly at –a*.125 and *a* if evidence initially crossed 0), and confidence was quantified at the time when the continued evidence accumulation crossed this second-order threshold. To project model confidence onto the same scale as human confidence, we used a linear transformation.

## Results

### Dynamic signatures of confidence

Following recent work that has modeled confidence as post-decision evidence accumulation (Calder-Travis et al., 2020; Pleskac & Busemeyer, 2010; Van Den Berg et al., 2016), we reasoned that confidence reports may differ, depending on whether they are probed around the time of the response (Kiani & Shadlen, 2009; Zylberberg et al., 2016) or only later in time, after additional post-decision processing (Moran et al., 2015; Pleskac & Busemeyer, 2010; Yu et al., 2015). In both cases, confidence reflects the probability of being correct, given the choice and accumulated evidence up until that point. The heat map in Fig. 1A reflects the probability of being correct given evidence (Y-axis) and time (X-axis), conditional on the choice (Kiani & Shadlen, 2009; Zylberberg et al., 2016). The current heat map shows this probability conditional on reaching the upper boundary (i.e., an “up” choice). Note that the heat map is flipped over the abscissa when the lower boundary is reached instead (i.e., a “down” choice). Thus, confidence in this model reflects the probability of being correct, given choice, evidence (potentially post-choice) and time. With respect to Signature 1 (an interaction between evidence strength and choice accuracy), initial model simulations show that confidence increases for both corrects and errors when confidence is quantified at the time the bound is reached (Figure 2B), whereas the interaction between evidence strength and choice accuracy emerges when confidence is queried after a small temporal delay (Figure 2C) and this interaction is most clearly seen after a more substantial period of post-decision processing (Figure 2D). In the following, we will test these predictions in human participants during random dot motion discrimination with additional confidence ratings.

### Perceptual decisions as noisy evidence accumulation

To unravel how coherence and volatility affected latent cognitive variables in the decision process, we fitted choices and reaction times using a hierarchical version of the drift diffusion framework (Wiecki et al., 2013). Because the effects of coherence and volatility were not modulated by the timing of confidence reports (immediate vs delayed) for both RTs, *F* < 1, *Bayes Factor (BF)* = .008, and accuracy, *F* < 1, *BF* = .01, the RT and accuracy data were combined. First, as typically observed in random dot motion tasks, drift rates increased monotonically with coherence level (see Figure 3A), with significant differences in drift rate between all coherence levels (averaged across volatility levels), *p*s < .001. Estimated drift rates did not depend on the level of evidence volatility, *p*s > .119. Second, as we predicted (Zylberberg et al., 2016), our manipulation of within-trial evidence volatility was captured by the within-trial drift variability parameter σ (see Figure 3B; Methods). When averaged over different coherences, estimated within-trial variability was higher for high compared to low volatility, *p* = .014 (pair-wise comparisons within each coherence value: 0% coherence: *p* = .091; 5% coherence: *p* = .049; 10% coherence: *p* = .259; 20% coherence: *p* = .106; 40% coherence: *p* = .457).

**Figure 3.**
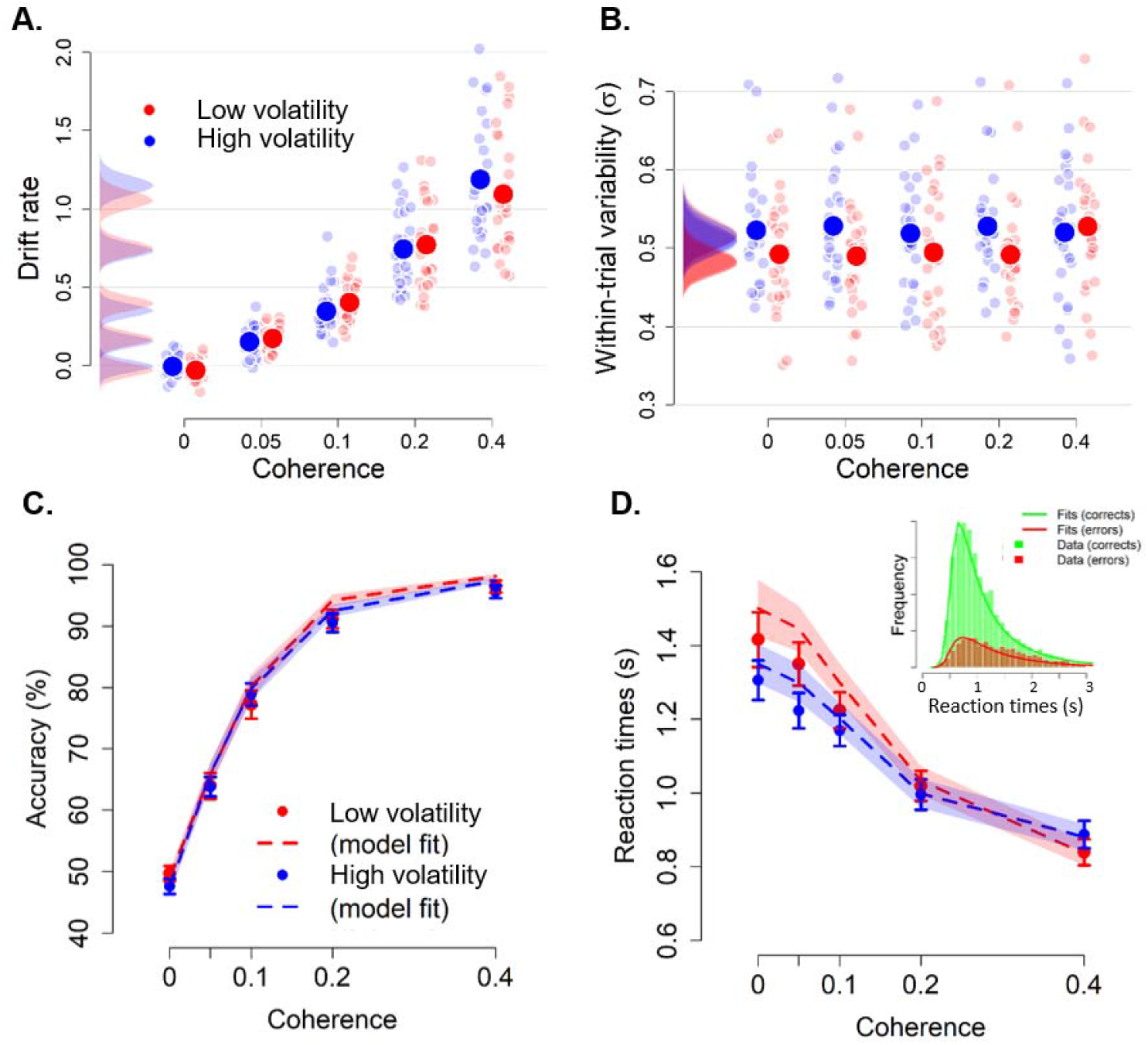
Model fits and task performance. **A**. Drift rate scales monotonically with the proportion of coherently moving dots, but did not differ for high and low volatility conditions. **B**. Within-trial variability (σ) selectively varied as a function of evidence volatility, whereas it was unaffected by motion coherence. Large dots: group averages; small dots: individual participants. Distributions show the group posteriors. Statistical significance is reflected in overlap between posterior distributions over parameter estimates (Materials and Methods). **C-D**. Accuracy (C) and RTs (D) as a function of coherence and evidence volatility, separately for the empirical data (points and bars) and model fits (lines and shades). Inset, distribution of empirical and fitted RTs. Shades and error bars reflect SEM of model and data, respectively.

These model fits captured the patterns seen in behavioral data (Figure 3C-D). Accuracy increased with the level of coherence (data: *F*(4,22) = 267.48, *p* < .001; model: *F*(4,22) = 619.57, *p* < .001), whereas evidence volatility and the interaction between both variables left accuracy unaffected (data: *F*s < 1; model: *p*s > .213). Reaction times decreased with increasing coherence levels (data: *F*(4,22) = 30.68, *p* < .001; model: *F*(4,22) = 52.25, *p* < .001), and were shorter with high compared to low volatility (data: *F*(1,25) = 9.10, *p* = .006; model: *F*(1,25) = 17.91, *p* < .001), an effect that was mostly pronounced at low coherence levels (data: *F*(4,22) = 13.21, *p* < .001; model: *F*(4,22) = 15.53, *p* < .001).

### Post-decision accumulation explains dynamic signatures of confidence

Next, we used our model fits to obtain qualitative and quantitative predictions of confidence reports about the three dynamic signatures of confidence. In order to create a heat map reflecting the probability of being correct, we simulated a large number of trials using the fitted drift rates and calculated average accuracy for each combination of time and evidence. Confidence predictions were quantified by reading out the values from this heat map (reflecting the probability of being correct) for each combination of evidence, time, and choice. The model predictions concerning confidence were highly similar when using an analytical solution (Moreno-Bote, 2010) instead.

#### Signature 1: interaction between evidence strength and choice accuracy

The first diagnostic signature of confidence established previously (Kepecs et al., 2008), is an increase of confidence with evidence strength for correct trials, but a decrease for error trials. In the immediate condition, confidence increased with coherence level, *F*(4,44.81) = 15.62, *p* < .001. Crucially, there was also the predicted interaction between coherence level and choice accuracy, *F*(4,1990.70) = 14.09, *p* < .001. Confidence increased with evidence strength for correct trials (linear contrast: *p* < .001), but there was no significant effect for error trials (linear contrast: *p* = .070; see Figure 4A). In contrast, as visualized in Figure 2B, the model predicts that when confidence is quantified at the time when the decision boundary is reached, confidence scales with coherence, *F*(4,25) = 32.76, *p* < .001, but there is no interaction between coherence and choice accuracy, *F* < 1.

**Figure 4.**
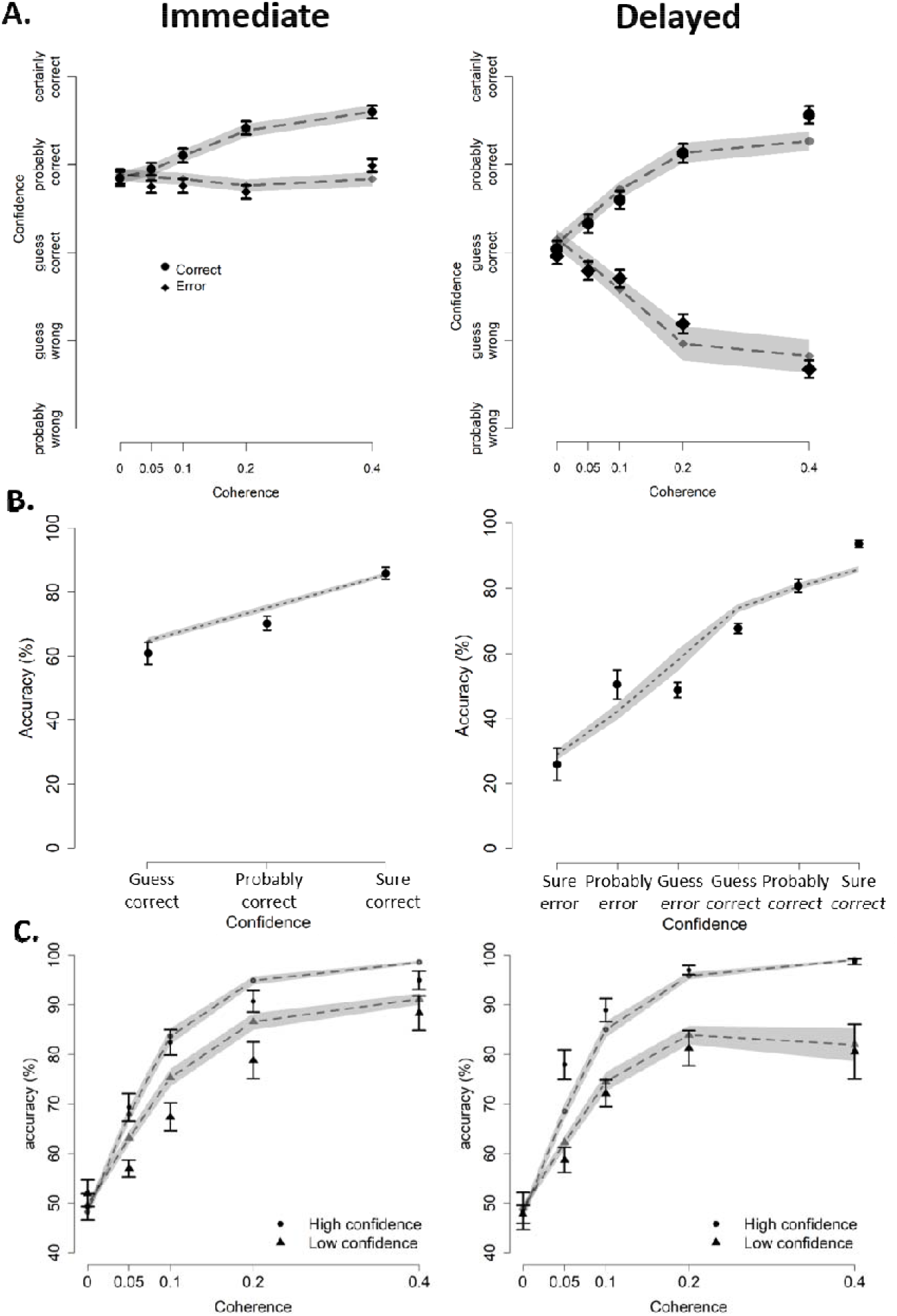
Three dynamic signatures of confidence. **A. Signature 1: an interaction between evidence strength and choice accuracy.** When confidence is quantified shortly after the decision bound has been reached (“immediate”), both model and data show an interaction between evidence strength and choice accuracy in the immediate condition. The same pattern was observed for the delayed condition, although the interaction effect was clearly much stronger here. **B. Signature 2: monotonically increasing accuracy as a function of confidence**. Both model and data show a monotonic scaling of accuracy depending on the level of confidence. **C. Signature 3: Steeper psychometric performance for high versus low confidence**. Both model and data show a steeper psychometric performance for trials judged with high versus low confidence. Notes: data for the delayed conditions are averaged over blank and extra evidence conditions. All plots show empirical data (black points and bars) and model predictions (grey lines and shades). Shades and error bars reflect SEM of model and data, respectively.

The above mismatch can easily be remedied by assuming that choice and confidence cannot be computed simultaneously, for example due to a brief refractory period (Marti et al., 2015; Pashler, 1994). Indeed, when confidence was calculated with a small temporal delay (100 ms, Figure 2C; see Methods), the model did predict the interaction between coherence and choice accuracy, *F*(4,200) = 153.09, *p* < .001. As in the behavioral data, the model with the small temporal delay predicted increasing confidence with coherence for correct trials (linear contrast: *p* < .001), but not for error trials (linear contrast: *p* = .541; Figure 4A). In the remainder, we will continue with predictions from the model with temporal delay.

In both delayed conditions, confidence scaled with coherence level (blank condition: *F*(4,51.8) = 5.49, *p* < .001; extra evidence condition: *F*(4,4571.1) = 4.75, *p* < .001). In both conditions, there was also an interaction between coherence and choice accuracy (blank condition: *F*(4,3625.6) = 53.38, *p* < .001; extra evidence condition: *F*(4,4568.7) = 71.45, *p* < .001). Within the correct trials, confidence increased with coherence levels (blank and extra evidence conditions, linear contrasts: *p* < .001. Instead, within the error trials, confidence decreased as a function of coherence (blank and extra evidence conditions, linear contrasts: *p* = .001). This interaction was captured by a model which terminated post-decision accumulation after a fixed amount of time (cf. Figure 2D; Materials and Methods). This model also showed the scaling of confidence with coherence (*F*(4,200) = 6.00, *p* < .001), as well as the interaction with choice accuracy (*F*(4,200) = 274.64, *p* < .001). Similar to the human data, confidence increased with coherence for correct trials (linear contrast: *p* < .001) and decreased for error trials (linear contrast: *p* < .001; figure 4B). Finally, there was a three-way interaction between coherence, choice accuracy and interrogation condition (data: *F*(8,13466.5) = 18.22, *p* < .001, model: *F*(4,450) = 181.88, *<* .001).

#### Signature 2: Monotonically increasing accuracy as a function of confidence

The second diagnostic signature of confidence is that it monotonically predicts choice accuracy. Indeed, an approximately linear relation between confidence and mean accuracy was observed in the data for both the immediate condition, *b* = .13, *t*(29.92) = 12.82, *p* < .001, the delayed blank, *b* = .12, *t*(27.29) = 16.12, *p* < .001, and the delayed extra evidence condition, *b* = .13, *t*(23.33) = 15.45, *p* < .001. This pattern was also captured by the model in the immediate condition, *b* = .10, *t*(24.9) = 22.75, *p* < .001, and in the delayed condition, *b* = .12, *t*(120.97) = 23.42, *p* < .001 (see Figure 4B). Note that these slopes did not differ depending on the moment in time when confidence was queried (data: X^2^ = 2.03, *p* = .363; model: X^2^ = 3.76, *p* = .152).

#### Signature 3: Steeper psychometric performance for high versus low confidence

The third diagnostic signature of confidence is that the relation between accuracy and evidence strength should be steeper for trials judged with high versus low confidence. The model predicts that this difference should be larger for the delayed compared to the immediate condition (Figure 4C). To test this prediction, confidence reports were divided into high or low using a split-median, separately per participant. As expected, the interaction between coherence and confidence in predicting accuracy was observed both in the immediate condition (data: *X*^*2*^(4) = 30.9, *p* < .001; model: *X*^*2*^(4) = 15.4, *p* < .001), and in the delayed condition (data: delayed blank: *X*^*2*^(4) = 84.15, *p* < .001, extra evidence: *X*^*2*^(4) = 56.64, *p* < .001; model: *X*^*2*^(4) = 3569.7, *p* < .001; see Figure 4C). Finally, there was a significant three-way interaction between coherence, confidence and interrogation condition (data: *X*^2^(8) = 24.51, *p* = .002; model: *X*^*2*^(4) = 62.11, *p* < .001).

### Evidence volatility dissociates time-based and evidence-based stopping criteria

If confidence is quantified after additional post-decision processing, a stopping rule has to be implemented determining at which point in time confidence is quantified. In the previous simulations, following previous research a time-based stopping rule was implemented (Pleskac & Busemeyer, 2010; Yu et al., 2015). Specifically, confidence was quantified after a specified amount of time has passed. An alternative implementation, however, is that the stopping rule for confidence reports is evidence-based, just like the stopping rule for the preceding choice process (Moran et al., 2015). According to this evidence-based stopping rule, after reaching the initial choice threshold, agents impose a second threshold and a delayed confidence report is given when this second threshold is reached. Because the static signatures discussed before do not arbitrate between the two delayed confidence stopping criteria (see Supplementary Materials), we next turn towards our manipulation of evidence volatility. Previous work has shown that an evidence-based model can explain the volatility effect on confidence for immediate confidence judgments (Zylberberg et al., 2016). According to Zylberberg et al. (2016) this is because noise pushes the evidence accumulation process faster toward the bound (i.e., it produces shorter reaction times) and shorter reaction times are associated with higher confidence (Kiani et al., 2014). As can be seen in Figure 5A, in the immediate condition our data did show a clear negative relation between confidence and reaction times on correct trials (data: *b* = −.19, *t*(24.2) = −4.89, *p* < .001; model: *b* = −.48, *t*(24.94) = −9.92, *p* < .001). In the delayed condition, a similar negative relation was apparent in the guess-correct to sure-correct range (see Figure 5B); but reaction times decreased again for confidence levels below guess-correct, i.e., for trials where participants identified their response as being an error. These probably reflect trials on which participants made fast errors which they then detected. This inverted U-shape was seen in both data and model (data: first-order polynomial: *b* = −18.01, *t*(7005) = −37.87, *p* < .001, second-order polynomial: *b* = −5.37, *t*(6999) = −11.69, *p* < .001; model: first-order polynomial: *b* = −84.21, *t*(2.006e+05) = −121.3, *p* < .001, second-order polynomial: *b* = −60.38, *t*(2.006e+05) = −88.8, *p* < .001). This finding again corroborates the notion that slower reaction times relate to lower confidence, a finding mostly expressed for immediate confidence. We reasoned that the same manipulation could be used to disentangle a time-based versus an evidence-based stopping rule for delayed confidence judgments.

**Figure 5.**
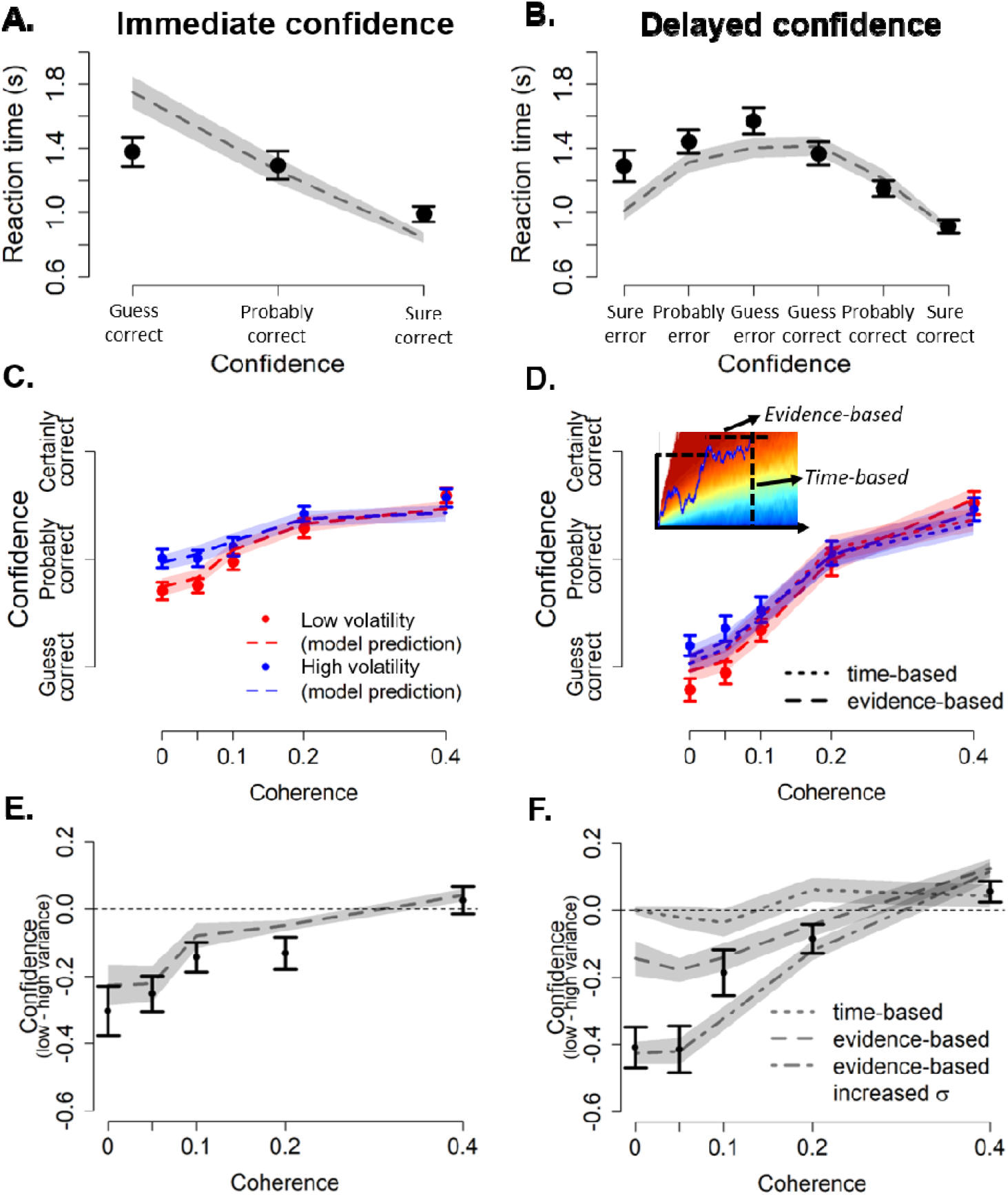
Within-trial evidence volatility arbitrates between an evidence-based and a time-based stopping rule. **A-B**. Average reaction times on correct trials as a function of decision confidence, separately for immediate (A) and delayed (B) confidence judgments. **C-F**. Immediate confidence (C and E) and delayed confidence (D and F) as a function of coherence and evidence volatility, separately for the empirical data (points and bars) and model predictions (lines and shades). C and D show average confidence, E and F show differences between low and high evidence volatility. The inset in D shows two potential stopping criteria for post-decision processing: post-decision accumulation can stop after a fixed period of time (i.e., a vertical time-based rule) or when a fixed amount of evidence is reached (i.e., a horizontal evidence-based rule). Notes: shades and error bars reflect SEM of model and data, respectively.

For *immediate confidence reports*, model predictions closely capture the pattern seen in human confidence ratings (see Figure 5C). Confidence monotonically increased with coherence levels (data: *F*(4,22) = 27.47, *p* < .001; model: *F*(4,22) = 27.68, *p* < .001), and was higher with high evidence volatility (data: *F*(1,25) = 41.19, *p* < .001; model: *F*(1,25) = 9.90, *p* = .004). Similar to RTs, the effect of evidence volatility on confidence was most pronounced with low coherence values (data: *F*(4,22) = 4.46, *p* = .008; model: *F*(4,22) = 30.79, *p* < .001). To easily interpret this effect, Figure 5E shows differences between the low and high volatility condition. As can be seen, for both model and data, confidence was increased with high evidence volatility, particularly with low coherence values.

For *delayed confidence* reports, the data favored the evidence-based stopping rule over the time-based stopping rule (see Figure 5D and 5F). The data and both models showed a monotonic increase of confidence with coherence levels (data extra evidence: *F*(4,22) = 46.67, *p* < .001; data blank: *F*(4,22) = 33.38, *p* < .001; time-based model: *F*(4,22) = 60.83, *p* < .001; evidence-based model: *F*(4,22) = 46.03, *p* < .001), and an interaction between coherence and volatility (data extra evidence: *F*(4,22) = 10.39, *p* < .001; data blank: *F*(4,22) = 8.42, *p* < .001; time-based model: *F*(4,22) = 11.94, *p* < .001; evidence-based model: *F*(4,22) = 23.50, *p* < .001). However, evidence volatility affected confidence in the data and the model with the evidence-based stopping rule (extra evidence: *F*(1,25) = 23.78, *p* < .001; blank: *F*(1,25) = 28.69, *p* < .001; evidence-based rule, *F*(1,25) = 8.96, *p* = .006), but not with the time-based stopping rule, *F* < 1. Finally, in the human data, delayed confidence reports were similar irrespective of whether post-decision evidence or a blank screen was presented following the choice (data not shown). This was further confirmed by an analysis including post-decision evidence (extra evidence or blank), which did not show a three-way interaction, *F* < 1, *BF* = .037.

Figure 5F suggests that the effect of volatility on confidence for the lowest coherence values is even stronger than predicted by the model with the evidence-based stopping rule. This is most likely because the sigma parameter, which captures evidence volatility, was estimated based on choices and RTs only (i.e., not based on confidence). Therefore, our predictions about immediate and delayed confidence are entirely constrained by the decision process itself. Some evidence hints at the possibility that post-decision accumulation is different from pre-decision accumulation (Yu et al., 2015). In the current context, it could therefore be that post-decision processing from memory amplifies noise in the sampling process. Indeed, when simulating the model with an evidence-based stopping rule using a slightly increased sigma value in the high volatility condition (σ = .575), it captures the pattern in the data even more tightly (see Figure 5F). This finding is in line with the possibility that post-decision accumulation is not fully determined by the pre-decision choice process.

## Discussion

Normative models explain the sense of confidence in a decision as the probability of a choice being correct. Although such formalization is principled and fruitful, it remains unclear whether and how it can account for dynamic expressions of confidence. To investigate this, we formalized confidence within an evidence accumulation framework as the probability of being correct, given the accumulated evidence up until that point. We tested model predictions concerning three diagnostic signatures of confidence, most notably an interaction between evidence strength and choice accuracy, both for immediate and delayed confidence reports. There was a close correspondence between model and human data for all three signatures, showing that these signatures of confidence depend on the time at which confidence is queried.

### Dynamic signatures of decision confidence

Static models have conceptualized confidence as the probability of being correct (Kepecs et al., 2008; Maniscalco & Lau, 2012; Sanders et al., 2016). Intuitively, when option A has a high (vs low) probability of being the correct answer, the model will give response A with high (vs low) confidence. One advantage of such a formalization is that it predicts three qualitative signatures of confidence (Sanders et al., 2016). A limitation of such an account is that this framework is inherently static, and therefore does not take time into account. To resolve this, we relied instead on a dynamic evidence accumulation framework to probe these different signatures across time. We are not the first to account for confidence within an evidence accumulation framework (Kiani et al., 2014; Moran et al., 2015; Pleskac & Busemeyer, 2010; Ratcliff & Starns, 2013; Zylberberg et al., 2016). Previous work has conceptualized immediate confidence as the probability of being correct given evidence and elapsed time (Kiani et al., 2014; Kiani & Shadlen, 2009; Zylberberg et al., 2016). Choices are formed when evidence reaches a fixed decision threshold, and both choice and confidence are quantified when this threshold is reached. As shown in Figure 2B, such a model does not predict an interaction between evidence strength and choice accuracy, a prediction at odds with many existing datasets. To account for this, we followed the approach taken by Pleskac and Busemeyer (2010) and allowed the evidence to continue accumulation following boundary crossing. By quantifying confidence across time, our model was able to account for these discrepancies. Specifically, our model was able to explain signature 1, an interaction between evidence strength and choice accuracy, in the immediate condition, as seen in behavioral data, by assuming that immediate confidence is quantified with a small temporal delay from the choice, suggesting a brief refractory period (Marti et al., 2015; Pashler, 1994). Thus, an important novel insight of the current work is that some form of post-decision evidence accumulation is necessary, even to explain immediate confidence reports. This also has implications for the way in which confidence judgments are elicited. Studies that query confidence only after the choice, on a scale which allows participants to indicate potential changes of mind (Boldt & Yeung, 2015; Van Den Berg et al., 2016), are more likely to find strong evidence for post-decision processing and more accurate confidence judgments compared to studies that simultaneously query choice and confidence (Kiani & Shadlen, 2009; Zylberberg et al., 2016). Our findings also have more widespread implications, for example with regard to literature on advice giving (Bonaccio & Dalal, 2006). Advice typically comes with a degree of confidence (Gaertig & Simmons, 2018). From the probabilistic perspective proposed here, this degree of confidence could also be taken into account when determining one’s own confidence, by conditioning on the confidence of the advice. The timing of such advice might be an important indicator of its accuracy, and thus its confidence. However, it remains an open question whether observers do indeed weight the advice of others depending on its timing.

As an important caution, we note that our conclusion about the necessity of post-decision accumulation might be limited to models with a static decision boundary. In models where the decision boundary collapses over the course of a trial, reaction times are predicted to be shorter for correct trials than for errors (J. Drugowitsch et al., 2012). Given the close link between reaction times and errors (i.e., as shown in Figure 5A), it remains a possibility that a model with a collapsing boundary is able to account for the interaction between evidence strength and choice correctness without post-decision evidence accumulation.

Previous modelling work has unraveled boundary conditions of this first diagnostic signature, the interaction between evidence strength and choice accuracy. Model simulations have shown that this interaction disappears if stimuli are only probabilistically related to choices (Adler & Ma, 2018), and if the static model has knowledge about evidence strength on the single-trial level (Rausch & Zehetleitner, 2019). Remarkably, however, no previous work has unraveled the role of time in this signature. The current work overcomes this limitation, by incorporating the notion of confidence reflecting the probability of being correct within a dynamic evidence accumulation framework. Our model simulations show that at the time of the boundary crossing, confidence increases with evidence strength for both corrects and errors, whereas the interaction effect only emerges with time. Crucially, this pattern was also observed in the empirical data. This has important consequences for studies relying on this signature to identify brain regions coding for decision confidence (Kepecs et al., 2008; Kepecs & Mainen, 2012). Because non-human animals cannot provide explicit confidence judgments, an alternative approach has been to test which brain regions show the three diagnostic signatures of confidence (Kepecs et al., 2008). Our results, however, suggest a slightly more complicated picture. Such regions should not show signature 1 (an interaction between evidence strength and choice correctness) at the time of choice commitment; they should, however, show this signature shortly after the time of commitment. In general, the magnitude of these three signatures in such brain regions should grow with post-decision time.

### Post-decision processing terminates using an evidence-based stopping rule

Post-decision evidence accumulation has been proposed as a mechanism explaining confidence (Moran et al., 2015; Pleskac & Busemeyer, 2010; Van Den Berg et al., 2016) and biases in confidence judgments (Navajas et al., 2016). It remains unclear, however, how such a model decides when to stop accumulating evidence. The decision process itself is believed to terminate once the accumulated evidence reaches a decision boundary. Likewise, our data favored an evidence-based stopping rule (i.e., the sampling process terminates when a certain level of evidence has been reached), while it was incompatible with a time-based stopping rule (i.e., sampling terminates after a certain time has elapsed). Only the evidence-based rule could explain increased confidence with high evidence volatility. Intuitively, high evidence volatility increases (immediate) confidence because the injection of noise in the decision process speeds up RTs (Zylberberg et al., 2016), and faster RTs are associated with higher confidence. The model with an evidence-based stopping rule for delayed confidence judgments similarly predicts higher confidence with high evidence volatility, because the noise again pushes the decision variable towards a certain level of evidence (i.e., a second bound). This effect does not appear with a time-based stopping rule, however, because the noise only affects the evidence (i.e., how fast is a certain level of evidence reached), but not the timing of post-decision accumulation itself. Therefore, using a time-based stopping rule the effects of evidence volatility are averaged out, and no differences in confidence are predicted. In sum, a second important insight of the current work is that human participants also use an evidence-based stopping rule in delayed confidence judgments.

### Sources of post-decisional evidence accumulation

The hypothesis that confidence is affected by post-decisional evidence accumulation has evoked a strong interest in neural signatures of post-decisional processing (Fleming et al., 2018; Murphy et al., 2015; Yu et al., 2015). For example, recent neuroimaging work has linked this process of post-decision evidence accumulation to a specific neural signal in the EEG (Murphy et al., 2015), that is sensitive to fine-grained levels of decision confidence (Boldt & Yeung, 2015; Desender, Murphy, et al., 2019). One question that has been largely overlooked so far, is what kind of information determines post-decisional evidence accumulation. For example, external information could drive post-decisional evidence accumulation (Fleming et al., 2018). Alternatively, internal sources, such as additional evidence from the sensory buffer (Resulaj et al., 2009) or resampling from memory (Vlassova & Pearson, 2013), could determine such accumulation. To contrast these two possibilities, the current work featured conditions with and without additional external evidence during the post-decisional period. Interestingly, confidence judgments were highly similar between these two conditions. This demonstrates that, at least in our current experimental design, participant benefit exclusively from internal resampling of the earlier evidence, whereas continued external sampling has no measurable influence. This does not imply that post-decisional evidence will never play a role in confidence. For example, in a recent study that de-correlated the strength of pre-decisional and post-decisional evidence (i.e., so that sometimes post-decision evidence was highly informative when pre-decision evidence was not), external post-decisional evidence did have a reliable effect on confidence (Fleming et al., 2018). Presumably, the correlational structure of post-versus pre-decision evidence determines whether sampling continues or not.

## Conclusion

The current work quantified confidence within an evidence accumulation framework as the probability of being correct given the accumulated evidence up until that point. Both model and data showed that three key signatures of confidence depend on the point in time when confidence is queried. Finally, post-decision confidence reports were best explained by an evidence-based stopping rule.

## Acknowledgments

The authors want to thank Peter R Murphy for advice on the modeling and Niklas Wilming, Cristian Buc Calderon and Konstantinos Tsetsos for fruitful discussions. This research was supported by an FWO [PEGASUS]^2^ Marie Sklodowska-Curie fellowship (12T9717N, to K.D.), DFG grants DO 1240/2-1 and DO 1240/3-1 (to. T.H.D.), and a Research Council-Flanders grant G010419N (to K.D and T.V.).

## Supplementary Materials

### Diagnostic confidence signatures with an evidence-based stopping rule

Model predictions about the diagnostic confidence signatures for the delayed condition were quantified using a time-based stopping rule. Here, we report that these predictions were highly similar when using an evidence-based stopping rule instead. First, this model also predicted that confidence scales with coherence, *F*(4,225) = 84.43, *p* < .001, as well as the interaction between coherence and choice accuracy, *F*(4,225) = 232.31, *p* < .001, reflecting increasing confidence with coherence levels for correct trials (linear contrast: *p* < .001) and decreasing for error trials (linear contrast: *p* < .001). Second, this model also predicted a monotonic positive relation between confidence and mean accuracy, *b* = .06, *t*(129) = 20.42, *p* < .001.

## Notes

### Competing Interest Statement

The authors have declared no competing interest.

## References

Adler, W. T., & Ma, W. J. (2018). Limitations of Proposed Signatures of Bayesian Confidence. Neural Computation, 30, 1–28. https://doi.org/10.1162/neco

Bates, D. M., Maechler, M., Bolker, B., & Walker, S. (2015). Fitting Linear Mixed-Effects Models Using lme4. Journal of Statistical Software, 67(1), 1–48.

Boldt, A., & Yeung, N. (2015). Shared Neural Markers of Decision Confidence and Error Detection. Journal of Neuroscience, 35(8), 3478–3484. https://doi.org/10.1523/JNEUROSCI.0797-14.2015

Bonaccio, S., & Dalal, R. S. (2006). Advice taking and decision-making: An integrative literature review, and implications for the organizational sciences. Organizational Behavior and Human Decision Processes, 101(2), 127–151. https://doi.org/10.1016/j.obhdp.2006.07.001

Brainard, D. H. (1997). The Psychophysics Toolbox. Spatial Vision, 10(4), 433–436. https://doi.org/10.1163/156856897X00357

Braun, A., Urai, A. E., & Donner, T. H. (2018). Adaptive history biases result from confidence-weighted accumulation of past choices. Journal of Neuroscience, 38(10), 2418–2429. https://doi.org/10.1523/JNEUROSCI.2189-17.2017

Calder-Travis, J., Bogacz, R., & Yeung, N. (2020). Bayesian confidence for drift diffusion observers in dynamic stimuli tasks. BioRxiv, 2020.02.25.965384. https://doi.org/10.1101/2020.02.25.965384

Desender, K., Boldt, A., Verguts, T., & Donner, T. H. (2019). Confidence predicts speed-accuracy tradeoff for subsequent decisions. ELife, e43499, p1:25. https://doi.org/10.1101/466730

Desender, K., Murphy, P., Boldt, A., Verguts, T., & Yeung, N. (2019). A Postdecisional Neural Marker of Confidence Predicts Information-Seeking in Decision-Making. The Journal of Neuroscience, 39(17), 3309–3319.

Drugowitsch, J., Moreno-Bote, R., Churchland, A. K., Shadlen, M. N., & Pouget, A. (2012). The Cost of Accumulating Evidence in Perceptual Decision Making. Journal of Neuroscience, 32(11), 3612–3628. https://doi.org/10.1523/JNEUROSCI.4010-11.2012

Drugowitsch, Jan, Moreno-Bote, R., & Pouget, A. (2014). Relation between belief and performance in perceptual decision making. PLoS ONE, 9(5). https://doi.org/10.1371/journal.pone.0096511

Fleming, S. M., van der Putten, E. J., & Daw, N. D. (2018). Neural mediators of changes of mind about perceptual decisions. Nature Neuroscience.https://doi.org/10.1038/s41593-018-0104-6

Gaertig, C., & Simmons, J. P. (2018). Do People Inherently Dislike Uncertain Advice? Psychological Science.https://doi.org/10.1177/0956797617739369

Gold, J. I., & Shadlen, M. N. (2007). The neural basis of decision making. Annual Review of Neuroscience, 30, 535–561. https://doi.org/10.1146/annurev.neuro.29.051605.113038

Kepecs, A., & Mainen, Z. F. (2012). A computational framework for the study of confidence in humans and animals. Philosophical Transactions of the Royal Society of London. Series B, Biological Sciences, 367(1594), 1322–1337. https://doi.org/10.1098/rstb.2012.0037

Kepecs, A., Uchida, N., Zariwala, H. a, & Mainen, Z. F. (2008). Neural correlates, computation and behavioural impact of decision confidence. Nature, 455(7210), p227–231. https://doi.org/10.1038/nature07200

Kiani, R., Churchland, A. K., & Shadlen, M. N. (2013). Integration of Direction Cues Is Invariant to the Temporal Gap between Them. Journal of Neuroscience, 33(42), p16483–16489. https://doi.org/10.1523/JNEUROSCI.2094-13.2013

Kiani, R., Corthell, L., & Shadlen, M. N. (2014). Choice Certainty Is Informed by Both Evidence and Decision Time. Neuron, 84(6), 1329–1342. https://doi.org/10.1016/j.neuron.2014.12.015

Kiani, R., & Shadlen, M. N. (2009). Representation of confidence associated with a decision by neurons in the parietal cortex. Science, 324(5928), 759–764. https://doi.org/10.1126/science.1169405

Kuznetsova, A., Brockhoff, P. B., & Christensen, R. H. B. (2014). lmerTest: Test in Linear Mixed Effect Models. R package version 2.0-20. http://CRAN.R-project.org/package=lmerTest.

Maniscalco, B., & Lau, H. (2012). A signal detection theoretic approach for estimating metacognitive sensitivity from confidence ratings. Consciousness and Cognition, 21(1), 422–430. https://doi.org/10.1016/j.concog.2011.09.021

Marti, S., King, J.-R., & Dehaene, S. (2015). Time-Resolved Decoding of Two Processing Chains during Dual-Task Interference. Neuron, 1–11. https://doi.org/10.1016/j.neuron.2015.10.040

Moran, R., Teodorescu, A. R., & Usher, M. (2015). Post choice information integration as a causal determinant of confidence: Novel data and a computational account. Cognitive Psychology, 78, 99–147. https://doi.org/10.1016/j.cogpsych.2015.01.002

Moreno-Bote, R. (2010). Decision confidence and uncertainty in diffusion models with partially correlated neuronal integrators. Neural Computation, 22(7), 1786–1811. https://doi.org/10.1162/neco.2010.12-08-930

Murphy, P. R., Robertson, I. H., Harty, S., & O’Connell, R. G. (2015). Neural evidence accumulation persists after choice to inform metacognitive judgments. ELife, 4(DECEMBER2015), 1–23. https://doi.org/10.7554/eLife.11946

Navajas, J., Bahrami, B., & Latham, P. E. (2016). Post-decisional accounts of biases in confidence. Current Opinion in Behavioral Sciences, 11, 55–60. https://doi.org/10.1016/j.cobeha.2016.05.005

Pashler, H. (1994). Dual-task interference in simple tasks: Data and theory. Psychological Bulletin, 116(2), 220–244. https://doi.org/10.1037/0033-2909.116.2.220

Pereira, M., Faivre, N., Iturrate, I., Wirthlin, M., Serafini, L., Martin, S., Desvachez, A., Blanke, O., de Ville, D. Van, & del Millán, J. R. (2020). Disentangling the origins of confidence in speeded perceptual judgments through multimodal imaging. Proceedings of the National Academy of Sciences of the United States of America, 117(15), 8382–8390. https://doi.org/10.1073/pnas.1918335117

Pleskac, T. J., & Busemeyer, J. R. (2010). Two-stage dynamic signal detection: A theory of choice, decision time, and confidence. Psychological Review, 117(3), 864–901. https://doi.org/10.1037/a0022399

Pouget, A., Drugowitsch, J., & Kepecs, A. (2016). Confidence and certainty: distinct probabilistic quantities for different goals. Nature Neuroscience, 19(3), 366–374. https://doi.org/10.1038/nn.4240

Ratcliff, R., & McKoon, G. (2008). The Diffusion Decision ModelD: Theory and Data for Two-Choice Decision Tasks. Neural Computation, 20, 873–922.

Ratcliff, R., & Starns, J. J. (2013). Modeling confidence judgments, response times, and multiple choices in decision making: recognition memory and motion discrimination. Psychological Review, 120(3), 697–719. https://doi.org/10.1037/a0033152

Rausch, M., & Zehetleitner, M. (2019). The folded X-pattern is not necessarily a statistical signature of decision confidence. PLoS Computational Biology, 15(10), e1007456. https://doi.org/10.1371/journal.pcbi.1007456

Resulaj, A., Kiani, R., Wolpert, D. M., & Shadlen, M. N. (2009). Changes of mind in decision-making. Nature, 461(7261), 263–266. https://doi.org/10.1038/nature08275

Sanders, J. I., Hangya, B., & Kepecs, A. (2016). Signatures of a Statistical Computation in the Human Sense of Confidence. Neuron, 90(3), 499–506. https://doi.org/10.1016/j.neuron.2016.03.025

Tuerlinckx, F., Maris, E., Ratcliff, R., & De Boeck, P. (2001). A comparison of four methods for simulating the diffusion process. Behavior Research Methods, Instruments, and Computers, 33(4), 443–456. https://doi.org/10.3758/BF03195402

Urai, A. E., Braun, A., & Donner, T. H. (2017). Pupil-linked arousal is driven by decision uncertainty and alters serial choice bias. Nature Communications, 8(14637), 1–11. https://doi.org/10.1038/ncomms14637

van den Berg, R., Zylberberg, A., Kiani, R., Shadlen, M. N., & Wolpert, D. M. (2016). Confidence Is the Bridge between Multi-stage Decisions. Current Biology, 26(23), p3157– 3168. https://doi.org/10.1016/j.cub.2016.10.021

Van Den Berg, R., Anandalingam, K., Zylberberg, A., Kiani, R., Shadlen, M. N., & Wolpert, D. M. (2016). A common mechanism underlies changes of mind about decisions and confidence. ELife, 1–21. https://doi.org/10.7554/eLife.12192

Vlassova, A., & Pearson, J. (2013). Look Before You Leap: Sensory Memory Improves Decision Making. Psychological Science, 24(9), 1635–1643. https://doi.org/10.1177/0956797612474321

Wiecki, T. V., Sofer, I., & Frank, M. J. (2013). HDDM: Hierarchical Bayesian estimation of the Drift-Diffusion Model in Python. Frontiers in Neuroinformatics, 7(August), 1–10. https://doi.org/10.3389/fninf.2013.00014

Yu, S., Pleskac, T. J., & Zeigenfuse, M. D. (2015). Dynamics of Postdecisional Processing of Confidence. Journal of Experimental Psychology: General, 144(2), 489–510.

Zylberberg, A., Fetsch, C. R., & Shadlen, M. N. (2016). The influence of evidence volatility on choice, reaction time and confidence in a perceptual decision. ELife, 5, 1–31. https://doi.org/10.7554/eLife.17688

